# *In vivo* genetic labeling of primary cilia in developing astrocytes

**DOI:** 10.1101/2025.03.19.644181

**Authors:** Rachel Bear, Claire Wei, Tamara Caspary

**Affiliations:** Department of Human Genetics, Emory University School of Medicine, 615 Michael Street Suite 301, Atlanta GA 30322, United States; Emory Graduate Program in Neuroscience

**Keywords:** Astrocytes, primary cilia, development, genetic models

## Abstract

Astrocyte cilia are largely understudied due to the lack of available tools. Astrocyte research advanced with the establishment of *Aldh1l1-Cre*^*ERT2*^, an inducible Cre line that specifically targets the astrocyte lineage. Here, we develop and compare genetic models that label astrocyte cilia in the developing prefrontal cortex (PFC) using *Aldh1l1-Cre*^*ERT2*^ and Cre-dependent cilia reporters. We evaluate these models by testing different tamoxifen-induction protocols and quantifying the percentage of astrocytes labeled with the cilia reporters. We show that tamoxifen dosage impacts the expression of cilia reporters in astrocytes. We validate the maximum cilia-labeling efficiency of tamoxifen using constitutively-expressed cilia reporters. The data reveal that only a subset of SOX9- positive astrocytes in the PFC possess cilia throughout development. Our work highlights the utility of Cre-Lox systems to target specific cell types and the importance of carefully validating genetic models.

## MAIN TEXT

Primary cilia are singular, antenna-like organelles that project from the cell membrane and are present in most vertebrate cell types. Cilia function as signaling centers (1). In the central nervous system (CNS), cilia are found on most neurons and glia and serve important functions during development (2). Astrocytes are the most abundant glial population and play critical roles in the development and function of the CNS (3). Cilia are required to regulate Sonic Hedgehog signaling (Shh) in astrocytes and function to modulate astrocyte development and homeostasis (4,5). However, our understanding of cilia function in astrocytes remains sparse due to the limited availability of tools to target and visualize astrocyte cilia *in vivo*.

Cilia localize many different signaling molecules that allow them to respond to extracellular signals and transduce these signals within the cell (6). In neurons, several signaling proteins localize specifically to cilia including adenylyl cyclase 3 (AC3) and somatostatin receptor 3 (SSTR3) (7,8). These ciliary proteins are predominantly expressed in neurons and are not present in most astrocytes (9,10). ADP-ribosylation factor-like protein 13B (ARL13B) is an atypical GTPase that influences ciliary structure and signaling (11). ARL13B is enriched along the ciliary membrane and is widely used as a cilia marker due to its nearly ubiquitous expression in primary cilia (12). ARL13B is expressed in ciliated cells throughout the CNS, including astrocytes (13). To identify astrocyte cilia, ARL13B staining requires co-staining with an astrocyte marker like SOX9, a nuclear protein expressed in nearly all astrocytes (14). The brain is densely populated with cilia such that it is challenging to distinguish which ARL13B-positive cilium is associated with which nucleus. This limits investigation of astrocyte cilia and fundamental questions remain about the frequency and composition of astrocyte cilia throughout the CNS.

Importantly, there are now Cre-Lox tools that permit genetic labeling of astrocyte cilia. First, the discovery of the ubiquitous astrocyte gene, aldehyde dehydrogenase 1 family member L1 (*Aldh1l1*), enables an astrocyte-specific Cre line (Cahoy et al., 2008). *Aldh1l1-Cre*^*ERT2*^ expresses tamoxifen-inducible Cre in over 90% of astrocytes in the CNS (Srinivasan et al., 2016). This overcomes the lack of specificity and astrocyte coverage of previous Cre lines (16). Second, there are Cre-dependent cilia reporters that express cilia-localized proteins tagged with fluorescent proteins, such as *Sstr3-Gfp* and *Arl13b-Cerulean* (18,19). These reporters label cilia regardless of whether the ciliary protein is normally expressed in that cell type. Here, we combine these tools to genetically label cilia in the astrocyte lineage.

To label cilia in the astrocyte lineage, we crossed *Aldh1l1-Cre*^*ERT2*^ mice to *Sstr3-Gfp* mice (hereafter referred to as *Sstr3-Gfp*^*Aldh1l1*^). We induced expression of *Sstr3-Gfp*^*Aldh1l1*^ by administering tamoxifen (TAM) using two different treatment protocols. Briefly, 2x TAM administers 2 doses of tamoxifen postnatally and 5x TAM administers 5 doses of tamoxifen embryonically (see methods and Figure 1a for details). We performed immunofluorescent staining for SOX9 and quantified astrocyte ciliation in the prefrontal cortex (PFC) at postnatal day (P) P8 and P21 (Figure 1b). In the 2x TAM treatment, we observed that 26.6% of astrocytes at P8 and 35.1% of astrocytes at P21 are ciliated in *Sstr3-Gfp*^*Aldh1l1*^ mice (Figure 1f). In the 5x TAM treatment, we observed that 61.4% of astrocytes at P8 and 64.7% of astrocytes at P21 are ciliated in *Sstr3-Gfp*^*Aldh1l1*^ mice (Figure 1g). This finding indicates that increased tamoxifen dosing in *Sstr3-Gfp*^*Aldh1l1*^ mice increases the labeling of astrocyte cilia. To examine whether 2x or 5x TAM is sufficient to label all astrocyte cilia, we generated *Sstr3-Gfp*^*ON*^ mice in which SSTR3-GFP is constitutively expressed in all cells (Figure 1c). We observed that 65.7% of astrocytes at P8 and 65.9% of astrocytes at P21 are ciliated in *Sstr3-Gfp*^*ON*^ mice indicating that 5x TAM treatment is sufficient to label all SSTR3-GFP-positive cilia in developing astrocytes (Figure 1h). These data suggest that a subset of astrocytes possess cilia.

**Figure 1:**
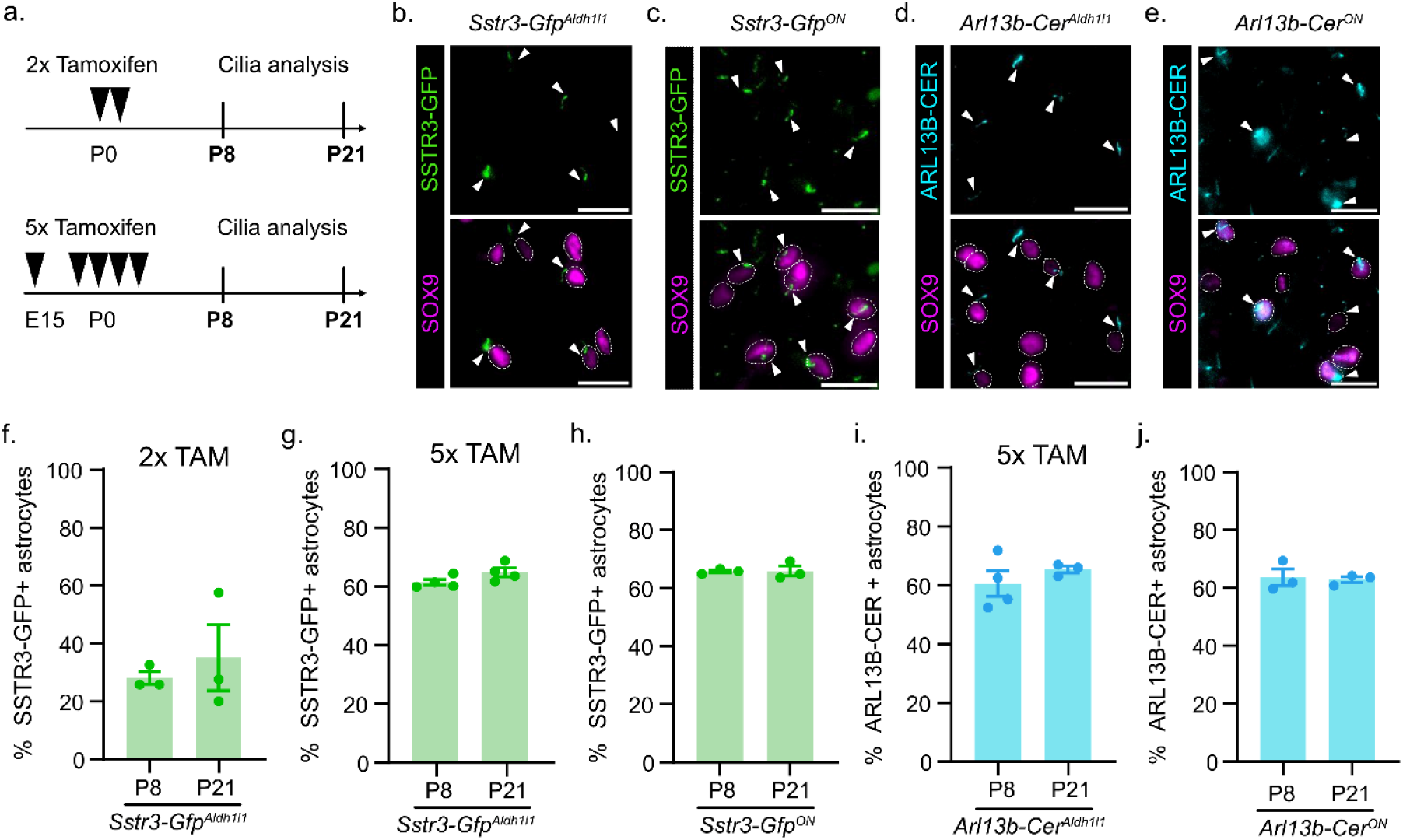
Expression of cilia reporters in the astrocyte lineage at P8 and P21. **(a)** Schematic of 2x Tamoxifen (TAM) and 5x TAM treatment protocol (black arrow) and astrocyte cilia analysis at P8 and P21. Representative images of labeled cilia (white arrow) in SOX9-positive astrocytes at P8 from **(b)** *Sstr3-Gfp*^*Aldh1l1*^ mice, **(c)** *Sstr3-Gfp*^*ON*^ mice, **(d)** *Arl13b-Cer*^*Aldlh1l1*^ mice, and **(e)** *Arl13b-Cer*^*ON*^ mice. Quantification of the percent SSTR3-GFP-positive cilia in SOX9-positive astrocytes in **(f)** *Sstr3-Gfp* ^*Aldlh1l1*^ mice using 2x TAM, **(g)** 5x TAM, and **(h)** *Sstr3-Gfp*^*ON*^ mice at P8 and P21. Quantification of the percent ARL13B-CER-positive cilia in SOX9-positive astrocytes in **(i)** *Arl13b-Cer* ^*Aldlh1l1*^ mice using 5x TAM and **(j)** *Arl13b-Cer* ^*ON*^ mice at P8 and P21. *n* = 3-4 animals. Scale bar = 20um. Data are expressed as the mean ± SEM.

To determine whether a different cilia reporter, *Arl13b-Cerulean*, functions equivalently, we incorporated it with the *Aldh1l1-Cre*^*ERT2*^ allele (hereafter referred to as *Arl13b-Cer*^*Aldh1l1*^). *Arl13b-Cerulean* and *Sstr3-Gfp* are both expressed at the *ROSA26* locus and contain a floxed STOP sequence for Cre-dependent activation. However, *Arl13b-Cerulean* is driven by a CAG promoter while *Sstr3-Gfp* uses the *ROSA26* promoter (18,19). To induce expression of *Arl13b-Cer*^*Aldh1l1*^, we administered 5x TAM (Figure 1a). We performed immunofluorescent staining for SOX9 and GFP to quantify the percentage of ciliated astrocytes (Figure 1d). We observed that 60.5% of astrocytes at P8 and 65.3% of astrocytes at P21 possess cilia (Figure 1i). This reveals that there is no difference in the percentage of astrocyte cilia labeled with 5x TAM in *Arl13b-Cer*^*Aldh1l1*^ compared to *Sstr3-Gfp*^*Aldh1l1*^ mice. To ensure 5x TAM is inducing complete expression of *Arl13b-Cer*^*Aldh1l1*^, we generated *Arl13b-Cer*^*ON*^ mice in which ARL13B-CER is expressed in all tissue (Figure 1e). We observed that 63.6% of astrocytes at P8 and 62.9% at P21 are ciliated in *Arl13b-Cer*^*ON*^ mice (Figure 1j). This result demonstrates that *Arl13b-Cer*^*ON*^ and *Sstr3-Gfp*^*ON*^ function equally to label astrocyte cilia. Furthermore, these data suggest that 5x TAM is sufficient to label all astrocyte cilia. At a biological level, these constitutive cilia labeling systems indicate that only a sub-population of astrocytes are ciliated and a non-ciliated sub-population also exists.

While there are many advantages for using the Cre-Lox system, the variability in recombination efficiency is a well-known limitation. First, the activity of the Cre line can vary in a tissue specific manner depending on the expression of the Cre driver (20). Second, there is high variation in recombination efficiency for different floxed alleles which can be attributed to the distance between the LoxP sites, sequences surrounding the LoxP sites, zygosity of the floxed allele, or the chromosomal location (20–22). Here, using the *Aldh1l1-Cre*^*ERT2*^ line, it is possible that *Aldh1l1* is not uniformly expressed among astrocytes and thus, may have a lower recombination efficiency. This could explain why tamoxifen dosage influenced the efficiency of cilia labeling in our model. Future studies are needed to tease apart whether the early administration and/or increased tamoxifen dosage is driving the increase of cilia reporter expression in astrocytes. We did not observe differences in the expression of the two cilia reporters, *Sstr3-Gfp* and *Arl13b-Cerulean*, suggesting differences in the reporter allele did not impact the recombination efficiency. These findings emphasize the importance of understanding Cre activity for specific floxed alleles in individual tissues.

In this study, we characterized several mouse models that label astrocyte cilia in the developing PFC. By achieving maximum labeling in our astrocyte cilia labeling system, we revealed that a sub-population of developing astrocytes are ciliated. This raises exciting questions about the specialized functions of the ciliated astrocyte population and whether there are brain region-specific differences. Our previous work demonstrates that astrocyte cilia are necessary for Shh signaling and that cilia influence astrocyte development in the PFC (4). Other questions include how the ciliated astrocyte population is established and whether they represent a distinct lineage from non-ciliated astrocytes. Thus, these genetic tools provide significant value to precisely target and label the ciliated astrocyte population.

## METHODS

### Mouse lines

All mice were cared for in accordance with NIH guidelines and Emory University’s Institutional Animal Care and Use Committee (IACUC). Lines used were *Aldh1l1-Cre*^*ERT2*^ obtained from The Jackson Laboratory [MGI:5806568, RRID:IMSR_JAX:031008] (16), *Sstr3-Gfp* obtained from Dr. Bradley Yoder [MGI: 5524281] (19), and *Arl13b-Cerulean* obtained through the European Mouse Mutant Archive [MGI:6193732, RRID:IMSR_EM:12168] (18). Genotyping was performed on DNA extracted from ear punch via PCR (36 cycles: 95°C 0.5min, 60°C 0.5min, 72°C 1min) using the primers listed (Table 1).

**Table 1:**
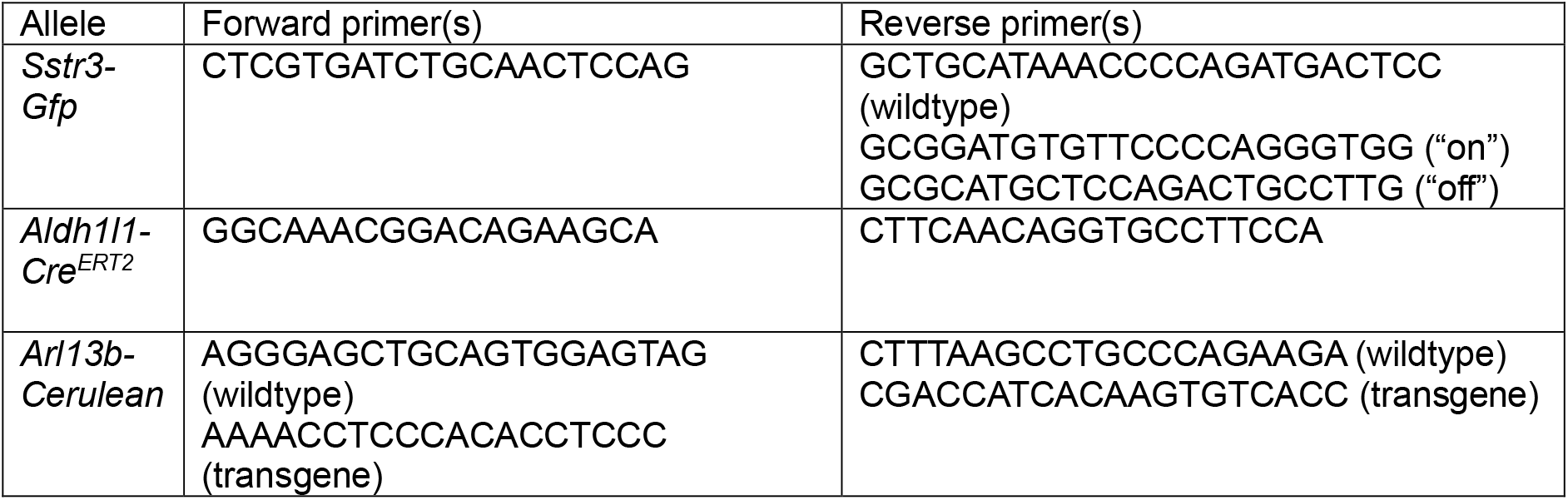
Genotyping Primers. Forward and reverse primer sequences used to genotype the mouse lines.

### Tamoxifen administration

Tamoxifen (Sigma T5648) stock solution was prepared once a month at a concentration of 10mg/ml in 100% EtOH and stored at −20°C. Each dose of tamoxifen was freshly prepared in corn oil the day of injection and dissolved using a speed vacuum centrifuge (Eppendorf Vacufuge Plus). To induce gene expression in astrocytes, a dose of 3mg tamoxifen/ 40g mouse weight in 300ul (adult) or 20ul (pup) of corn oil was prepared. For 2x tamoxifen treatment, tamoxifen was administered intraperitoneally at P0 and P1 to the dam using a 1ml syringe and a 25G 5/8-inch needle. For 5x tamoxifen treatment, tamoxifen was administered intraperitoneally at E15.5 and E19.5 to the pregnant dam using a 1ml syringe and a 25G 5/8-inch needle followed by subcutaneous administration at P0, P1, and P2 to pups using a 1/2ml syringe with attached 29G 1/2-inch needle.

### Caesarean section and cross fostering

A Caesarean section was performed on tamoxifen-treated, timed-pregnant dams at E20.5. CD-1 mice were used as foster dams. Briefly, the experimental dams were euthanized via cervical dislocation and the pups were dissected, placed on a heating pad, and tapped gently until they could breathe on their own. Pups were transferred to a CD-1 foster dam and integrated with the foster dam’s litter.

### Tissue harvesting

P21 mice were euthanized by isoflurane inhalation followed by a trans-cardiac perfusion with ice-cold 1x phosphate-buffered saline (PBS) and ice-cold 4% paraformaldehyde (PFA). P8 pups were euthanized by decapitation. Brains were harvested following perfusion or decapitation and drop-fixed in 4% PFA overnight. Fixed tissue was washed with 1x PBS and then incubated with 30% sucrose in 0.1 M phosphate buffer overnight at 4°C until the tissue sank. Samples were washed in optimal cutting temperature (OCT) compound to remove sucrose (Tissue-Tek OCT, Sakura Finetek), embedded in OCT, frozen on dry ice, and stored at −20°C.

### Immunofluorescent (IF) staining

OCT-embedded tissues were sectioned at 40µm using a cryostat microtome and placed directly on microscope slides (Fisherbrand Superfrost Plus). Sections were let to air-dry and either processed immediately or stored at −20°C. Sections stored at −20°C were brought to room temperature before starting IF. Sections were first rehydrated in 1X Tris Buffered Saline (TBS), permeabilized in 1% SDS, and blocked in antibody wash (1% heat inactivated goat serum, 0.1% Triton X-100 in 1X TBS). Sections were incubated overnight at 4°C with primary antibodies (chicken anti-GFP, 1:8,000, Abcam ab13970; rabbit anti-SOX9, 1:500, Millipore AB5535). Sections were washed with cold antibody wash and incubated for one hour at room temperature in the dark with secondary antibodies (goat anti-chicken, AlexaFluor 488; goat anti-rabbit, AlexaFluor 647; Hoechst 33342; all 1:500 dilution, ThermoFisher). Sections were washed with cold antibody wash and then mounted with a glass coverslip using ProLong Gold (ThermoFisher) mounting media. Slides cured overnight at room temperature in the dark and were stored short term at 4°C or long term at −20°C. Slides were imaged at 20x on a BioTek Lionheart FX automated microscope.

### Quantification of cilia

Z-stack images of brain sections stained for ciliary markers were captured at 2um intervals. Analysis of cilia was performed on maximum projection images using cell counting in Fiji/ImageJ.

## DECLARATIONS

### Ethics approval and consent to participate

All methods were carried out in accordance with guidelines and regulations of the National Institutes of Health guidelines and Emory University Institutional Animal Care and Use Committee for animal research. All experimental protocols using animals were approved by Emory University’s Institutional Animal Care and Use Committee.

### Consent for publication

Not applicable

### Availability of data and materials

The datasets used and/or analyzed during the current study are available from the corresponding author on reasonable request.

### Competing interests

The authors declare no competing interests.

### Funding

This work was supported by the National Institutes of Health T32NS096050 and F31NS125984 to RB, and R35GM122549 and R35GM148416 to TC.

### Author’s contributions

Conceptualization: RB and TC, Investigation: RB and CW, Methodology: RB and TC, Visualization: RB, Writing-original draft: RB, Writing-review and editing: RB, CW, and TC.

## Notes

### Competing Interest Statement

The authors have declared no competing interest.

